# *period* Translation as a Core Mechanism Controlling Temperature Compensation in an Animal Circadian Clock

**DOI:** 10.1101/2022.02.21.481387

**Authors:** Taichi Q. Itoh, Evrim Yildirim, Satya Surabhi, Rosemary Braun, Ravi Allada

## Abstract

The most enigmatic of the canonical properties of circadian clocks is temperature compensation where circadian period length is stable across a wide temperature range despite the temperature dependence of most biochemical reactions. While the core mechanisms of circadian clocks have been well described, the molecular mechanisms of temperature compensation are poorly understood especially in animals. A major gap is the lack of temperature compensation mutants that do not themselves unambiguously affect the temperature dependence of the encoded protein. Here we show that null alleles of two genes encoding components of a complex important for translation of the core clock component *period* in circadian pacemaker neurons robustly alter the temperature dependence of circadian behavioral period length. These changes are accompanied by parallel temperature dependent changes in oscillations of the PER protein and are consistent with the model that these translation factors mediate the temperature-dependence of PER translation. Consistent with findings from modeling studies, we find that translation of the key negative feedback factor PER plays an instrumental role in temperature compensation.

## Introduction

Circadian clocks have evolved to appropriately align the timing of behavior and physiology across a wide range of naturally occurring environmental conditions(1, 2). These clocks drive daily rhythms that free run under constant conditions, exhibit a periodicity around, but not exactly, 24 h, and are synchronized or entrained to the 24 h environment by cues such as light and temperature. Despite the sensitivity of the clock to changes in temperature, circadian period length remains relatively constant across a wide physiological temperature range, a phenomenon known as temperature compensation. How the circadian clock compensates for temperature remains largely unknown.

While most biochemical reactions significantly accelerate with increasing temperature (Q_10_ ∼2-3, i.e. rates increase 2-3x with a 10°C temperature increase), circadian period remains relatively constant (Q_10_∼0.8-1.2)(3). Temperature compensation has been observed in a wide range of organisms including cyanobacteria(4), *Neurospora*(5), *Arabidopsis*(6), in vitro clocks(7), and even in explanted tissues and cells from homeotherms(8-12), indicating that it is an intrinsic property of independently evolved clocks likely important for organismal fitness. Interestingly, circadian rhythms driven by coupled pacemaker neurons, e.g. the suprachiasmatic nucleus in mammals or the lateral neurons in *Drosophila*, exhibit even more precise Q_10_ (0.96-1.06)(3, 13, 14) suggesting a role for oscillator coupling in temperature compensation.

A variety of models have been proposed to explain this phenomenon. Hastings and Sweeney(15) proposed the model of balanced reactions in which the circadian clock consists of temperature-dependent reactions with opposing effects (e.g., inhibition of an activator) to maintain a near constant period. Balanced models have been generated that can recapitulate the temperature compensation of circadian rhythms (16), although this idea has been challenged(17, 18). While dozens of circadian mutants have been identified that alter circadian period, surprisingly, most remain temperature compensated. This observation runs counter to the prediction of the balanced model where perturbation of one temperature dependent step of the clock might reveal the temperature dependence of the remaining steps. An alternative is that there is a relatively discrete temperature dependent feedback mechanism within an otherwise temperature-independent clockwork that maintains period length. Temperature-dependent changes in amplitude have been proposed to play a role(18, 19). Nonetheless, the components of this mechanism remain to be elucidated.

In animals, circadian behavioral rhythms are driven by molecular feedback loops with transcriptional, posttranscriptional, and posttranslational regulatory layers(2, 20, 21). In flies, the heterodimeric transcription factor CLOCK (CLK)/CYCLE (CYC) activates *period (per)* and *timeless (tim)* transcription. PER and TIM proteins repress CLK-CYC. PER and TIM phosphorylation regulates their localization, function, and stability. These molecular oscillators are evident in both the central pacemaker neurons that govern circadian behavior as well as in peripheral tissues. A network of coupled pacemakers with the small ventral lateral neurons (sLNv) acting as the dominant pacemaker plays a critical role in timing free-running behavioral rhythms(22). Notably, these networked pacemakers employ an additional regulatory step critical for rhythmicity: translational activation of *per* mRNA via an RNA-binding complex of TWENTYFOUR (TYF), ATAXIN2 (ATX2), and LSM12(23-25). Cell autonomous molecular clocks collaborate with coupling of networked oscillators to produce robust free-running circadian rhythms(22).

A major concern with the existing mutants that alter temperature compensation in *Drosophila* (26-30) and many of those in other organisms(31-33), is that they are due to missense mutants that may alter the Q_10_ of a protein important for a rate-limiting step in setting circadian period. Thus, temperature compensation phenotypes may simply be due to temperature dependent misfolding, for example, rather than an intrinsic compensation phenotype. Indeed, about 10% of missense mutations in the p53 gene (as an example) exhibit temperature sensitive phenotypes(34), a rate that roughly approximates the proportion of missense clock mutants that have temperature compensation phenotypes, consistent with this hypothesis. The more definitive identification of *bona fide* temperature compensation mutants employs changing gene dosage or otherwise manipulating protein function without altering the Q_10_ of the encoded protein. This was used successfully in fungi to define a role for the protein kinase CK2 in altering temperature compensation, suggesting a role for phosphorylation(5) as has been proposed in mammalian clocks(31). Although even in this case some of the CK2 mutants actually exhibit better temperature compensation, i.e., a Q_10_ even closer to 1, than wild-type controls(5). In more precisely compensated animal clocks, inhibition of the steeply temperature sensitive heat shock transcription factor strongly alters the Q_10_ of molecular oscillations in explanted suprachiasmatic nuclei(12). Here we show that near null insertional alleles of the *per* translation factors *tyf* and *Lsm12* independently exhibit robust temperature compensation phenotypes, providing novel insights into the mechanistic basis of temperature compensation in animals and its link to PER translation.

## Results

To address the role of *per* translation in temperature compensation, we examined circadian period length in flies lacking the *per* translation activator *tyf*. As *tyf* mutants display poor circadian rhythms, it is difficult to assess circadian period length(25). However, we previously demonstrated that additional copies of a 13.2 kb genomic *per* transgene (*per13*.*2*) can partially rescue circadian behavioral rhythms enabling an assessment of circadian rhythms(25). To assess temperature compensation, we assessed circadian behavior at 18°C, 21.5°C, 25°C and 28°C, plotted temperature versus period length, and computed the Q_10_ (see Methods). As period estimation in flies with poor rhythms can be imprecise, we also separately analyzed those flies with robust rhythms, i.e, chi-square periodogram power-significance >40. Consistent with prior studies, we observed robust temperature compensation in a wild-type *iso31* strain (Q_10_=1.02; Table 1; Figure 1A,B). To determine if the *per13*.*2* transgene exhibited robust temperature compensation on its own we examined wild-type flies bearing two *per13*.*2* transgenes as well as *per*^*01*^ flies in which rhythms were rescued by *per13*.*2*. In each case, these flies exhibited robust temperature compensation (*per13*.*2* Q_10_=1.02, *per*^*01*^; *per13*.*2* Q_10_=0.99; Table 1, Figure 1A, B).

**Table 1.** Q10 of behavioral period in *tyf* mutants

| Genotype | Temperature<br>(°C) | N | Power +/- SEM | P-S>10 |  |  |  |  | P-S>40 |  |  |  |  |
| --- | --- | --- | --- | --- | --- | --- | --- | --- | --- | --- | --- | --- | --- |
|  |  |  |  | N | Period +/- SEM | Q10 | 2.5%<br>CI | 97.5%<br>CI | N | Period +/- SEM | Q10 | 2.5%<br>CI | 97.5%<br>CI |
| iso31 | 18 | 26 | 57 +/- 8 | 23 | 23.8 +/- 0.1 | 1.02 | 1.01 | 1.03 | 16 | 23.7 +/- 0.1 | 1.02 | 1.01 | 1.03 |
|  | 21.5 | 15 | 49 +/- 8 | 14 | 24.0 +/- 0.1 |  |  |  | 13 | 23.9 +/- 0.1 |  |  |  |
|  | 25 | 10 | 79 +/- 14<sup>b</sup> | 9 | 23.7 +/- 0.1 |  |  |  | 8 | 23.8 +/- 0.1 |  |  |  |
|  | 28 | 25 | 89 +/- 8<sup>a,b</sup> | 24 | 23.3 +/- 0.1<sup>a,b</sup> |  |  |  | 23 | 23.3 +/- 0.1<sup>a,b,c</sup> |  |  |  |
| per<sup>0</sup>; per13.2 | 18 | 29 | 29 +/- 7 | 15 | 23.2 +/- 0.2 | 0.98* | 0.97 | 1.00 | 8 | 22.2 +/- 0.1 | 0.97* | 0.96 | 0.99 |
|  | 21.5 | 23 | 53 +/- 8 | 20 | 23.4 +/- 0.1 |  |  |  | 16 | 23.3 +/- 0.1 |  |  |  |
|  | 25 | 24 | 80 +/- 10<sup>a</sup> | 22 | 23.6 +/- 0.1 |  |  |  | 18 | 23.6 +/- 0.1<sup>a</sup> |  |  |  |
|  | 28 | 16 | 58 +/- 12 | 12 | 23.5 +/- 0.1 |  |  |  | 19 | 23.5 +/- 0.1<sup>a</sup> |  |  |  |
| per13.2 | 18 | 67 | 56 +/- 5 | 52 | 23.1 +/- 0.1 | 1.02 | 1.01 | 1.03 | 40 | 23.2 +/- 0.1 | 1.02 | 1.01 | 1.03 |
|  | 21.5 | 44 | 65 +/- 6 | 39 | 23.0 +/- 0.1 |  |  |  | 33 | 23.0 +/- 0.1 |  |  |  |
|  | 25 | 32 | 68 +/- 10 | 24 | 22.9 +/- 0.1 |  |  |  | 20 | 23.0 +/- 0.1 |  |  |  |
|  | 28 | 44 | 116 +/- 7<sup>a,b,c</sup> | 43 | 22.7 +/- 0.0<sup>a,b</sup> |  |  |  | 41 | 22.7 +/- 0.0<sup>a,b,c</sup> |  |  |  |
| tyf<sup>e</sup> | 18 | 70 | 3 +/- 1 | 6 | 25.4 +/- 1.1 | 0.98 | 0.93 | 1.03 | 0 | - | 0.90* | 0.78 | 1.05 |
|  | 21.5 | 26 | 18 +/- 6 | 11 | 25.2 +/- 0.3 |  |  |  | 3 | 25.0 +/- 0.3 |  |  |  |
|  | 25 | 24 | 21 +/- 6<sup>a</sup> | 11 | 26.3 +/- 0.2 |  |  |  | 5 | 25.9 +/- 0.3 |  |  |  |
|  | 28 | 20 | 6 +/- 2<sup>c</sup> | 5 | 25.1 +/- 0.2 |  |  |  | 0 | - |  |  |  |
| tyf<sup>e</sup>;per13.2/+ | 18 | 30 | 11 +/- 4 | 9 | 22.6 +/- 0.2 | 0.80* | 0.78 | 0.83 | 3 | 22.8 +/- 0.3 | 0.80* | 0.78 | 0.83 |
|  | 21.5 | 24 | 55 +/- 7<sup>a</sup> | 22 | 23.8 +/- 0.1<sup>a</sup> |  |  |  | 16 | 23.8 +/- 0.1<sup>a</sup> |  |  |  |
|  | 25 | 52 | 32 +/- 5<sup>a,b</sup> | 35 | 26.1 +/- 0.1<sup>a,b</sup> |  |  |  | 20 | 26.1 +/- 0.1<sup>a,b</sup> |  |  |  |
|  | 28 | 24 | 11 +/- 4<sup>c</sup> | 7 | 27.9 +/- 0.9<sup>a,b,c</sup> |  |  |  | 1 | 28.0<sup>a,b,c</sup> |  |  |  |
| tyf<sup>e</sup>;per13.2 | 18 | 60 | 7 +/- 3 | 7 | 21.7 +/- 0.2 | 0.80* | 0.79 | 0.82 | 3 | 21.7 +/- 0.3 | 0.81* | 0.78 | 0.83 |
|  | 21.5 | 17 | 44 +/- 10<sup>a</sup> | 13 | 23.3 +/- 0.1<sup>a</sup> |  |  |  | 22 | 23.4 +/- 0.1<sup>a</sup> |  |  |  |
|  | 25 | 38 | 49 +/- 6<sup>a</sup> | 31 | 24.4 +/- 0.1<sup>a,b</sup> |  |  |  | 20 | 24.4 +/- 0.1<sup>a,b</sup> |  |  |  |
|  | 28 | 16 | 36 +/- 9<sup>a</sup> | 7 | 27.8 +/- 0.3<sup>a,b,c</sup> |  |  |  | 6 | 27.8 +/- 0.4<sup>a,b,c</sup> |  |  |  |

Statistical significances in period and power were determined using one-way ANOVA followed Tukey's multiple comparison test. a, b, and c indicate significantly difference with 18°C, 21.5°C and 25°C, respectively. Asterisks show statistically difference between *iso31* and each genotype. CI indicates confidence intervals.

**Figure 1.**
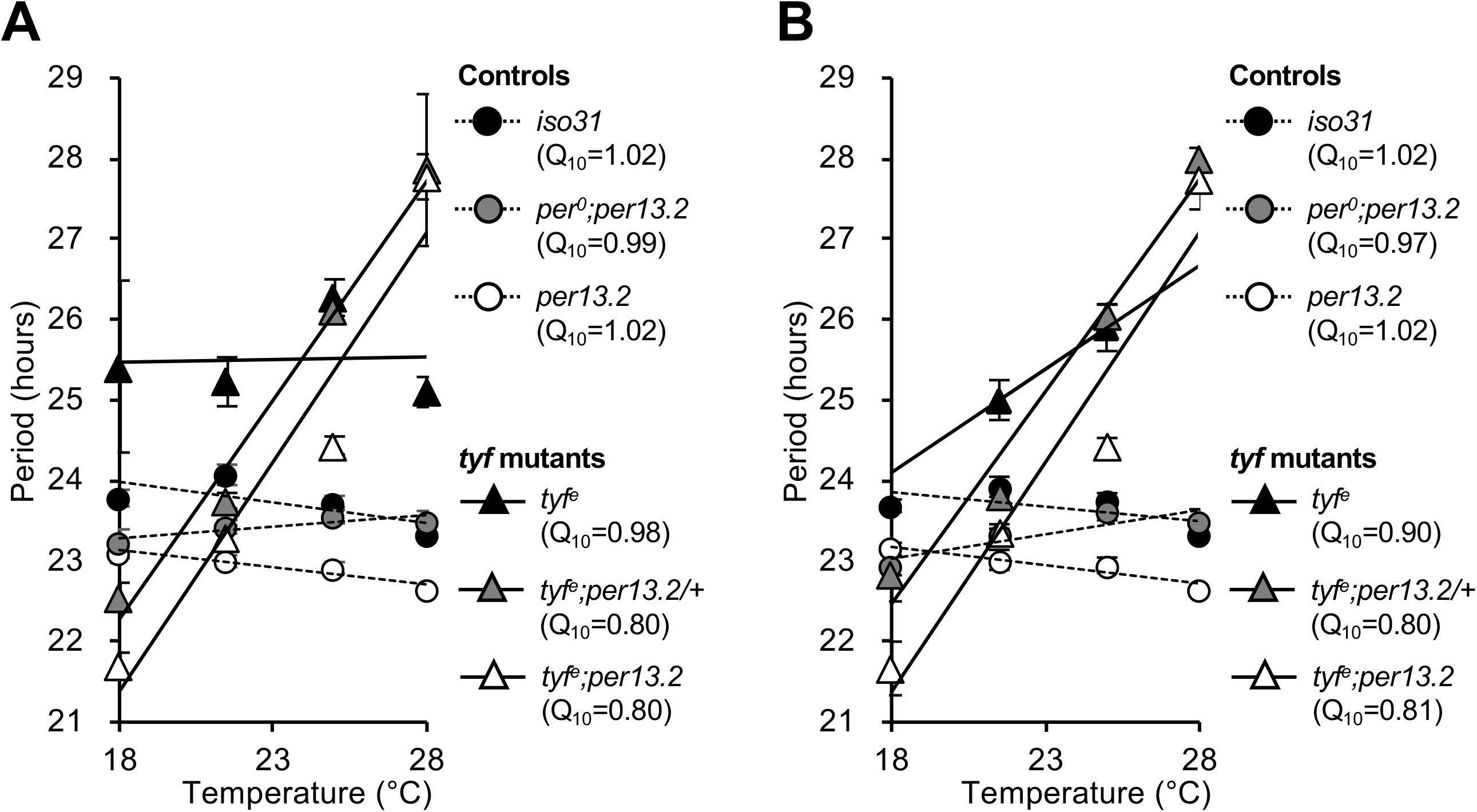
Lack of TYF induces a substantial defect in temperature compensation. Averaged periods of the free-running periods under four different temperatures were calculated only from the flies showing power - significance values (P-S) over 10 (A) and 40 (B) at the corresponding temperatures. Data represents average +/-SEM. The dashed and solid lines represent their regression lines of controls and tyf mutants, respectively. *tyf*^*e*^ line is a null mutant of *tyf*, and *per13*.*2* line is carrying a *period* genomic construct on 3rd chromosome. Sample sizes are indicated in Table 1.

As expected, *tyf* mutants displayed poor rhythmicity with no flies exhibiting robust rhythms (P-S>40) at 18 °C or 28 °C, making it difficult to robustly assess temperature compensation. Of note, the few robustly rhythmic *tyf* flies displayed a modest (∼1h) but significant (p<10^−10^) period lengthening between 21.5°C and 25°C(Table 1, Figure 1A, B). To more rigorously assess temperature compensation in a *tyf* mutant background, we added *per13*.*2* transgenes which dramatically improved rhythmicity enabling more precise period estimates. Here we observed strong temperature compensation phenotypes with period lengths ∼22h at 18°C and ∼27h at 28°C (Q_10_=0.80 with either one or two *per13*.*2* transgenes; Table 1, Figure 1A, B). Thus, we demonstrate that *tyf* is required for robust temperature compensation without altering the Q_10_ of any individual gene/protein.

To assess temperature dependence in *tyf* mutants without *per13*.*2*, we assessed their behavior phase on days 1 and 2 of DD. As *tyf* mutants display intact evening anticipation under LD conditions(25), we reasoned that DD rhythms may be more clearly evident early in DD and then damp over time. Examining evening phase offset at 18°C on the first and second days of DD, we found that *tyf* mutants are phase advanced on the first (CT12.24 v. CT10.07; p<0.05) and second (CT36.08 v. CT33.22; p<0.05; Figure 2; Table 2). However at 28°C, we find that *tyf* mutants are phase delayed relative to wild-type (DD1 CT15.04 v CT17.75; p<0.05, DD2 CT38.80 v. CT44.49; p<0.05; Figure 2; Table 2), consistent with a temperature-dependent period lengthening. A similar phenotype was observed in *tyf; per13*.*2* flies. This temperature dependence was evident early in DD with ∼3h phase advances relative to wild type observed at 18°C (DD1/2: CT12.31/CT35.06 v. CT9.53/C31.36, p<0.05; Figure 2; Table 2) consistent with a shorter period and ∼6h (on DD2) phase delays at 28°C (DD1/2: CT14.90/CT37.36 v. CT16.14/CT43.52, p<0.05; Figure 2; Table 2), consistent with a longer period in *tyf; per13*.*2* flies.

**Figure 2.**
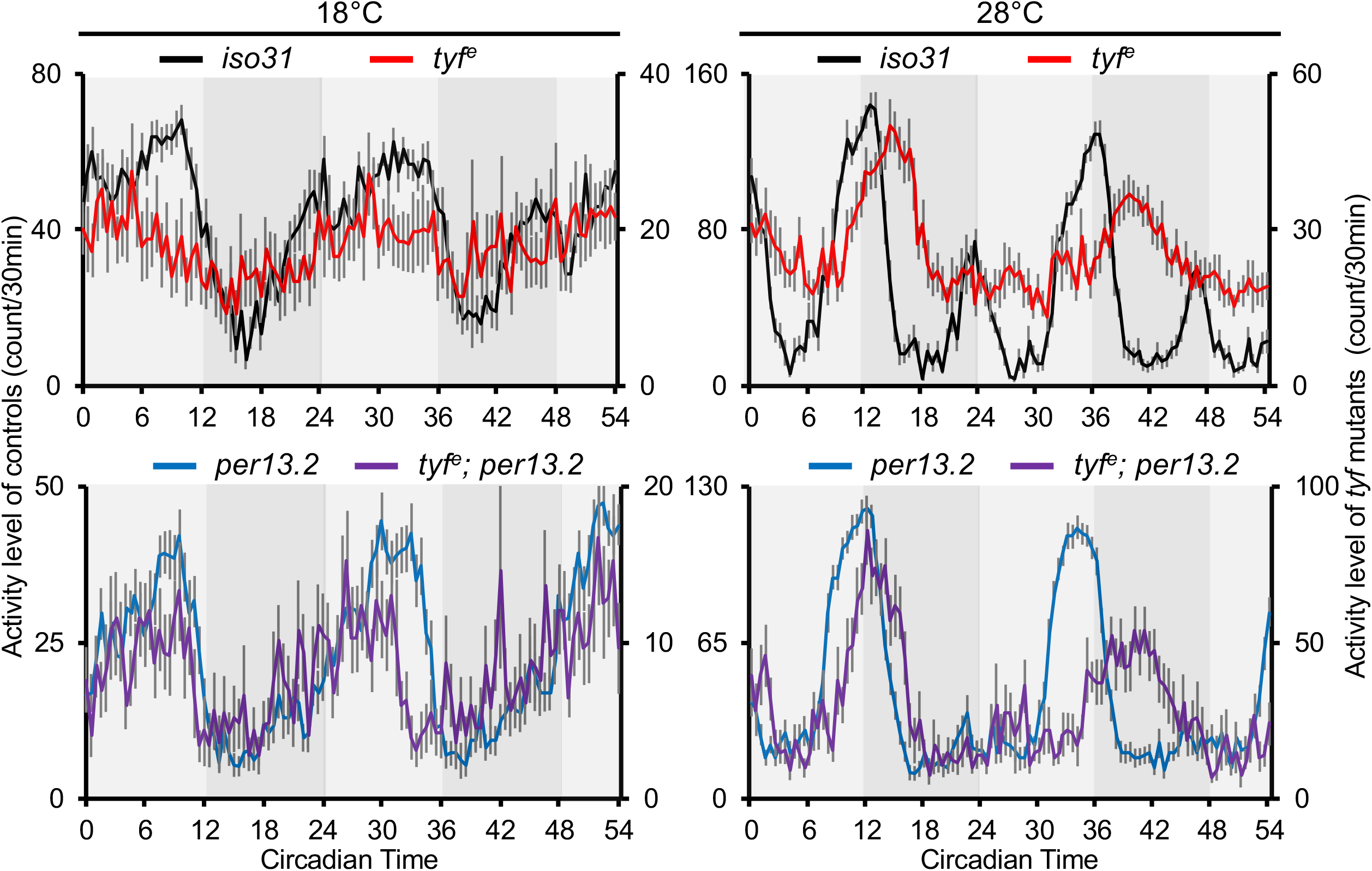
Peak phases of activity level in tyf mutants advance in 18C and delay in 28C. Data represents averaged activity levels +/-SEM each 30 min from CT0 to CT54. First axis indicates activity level of controls, and the second axis is that of *tyf* mutants or *tyf* mutants with *per* genomic rescue. Color codes and sample sizes are as follows. Black: *iso31* (n=27 at 18 and n=26 at 28°C), Red: *tyf*^*e*^ (n=69 at 18°C and n=25 at 28°C). Blue: *per13*.*2* (n=67 at 18°C and n=42 at 28°C), Purple: t*yf*^*e*^*;per13*.*2* (n=44 at 18°C and n=14 at 28°C).

**Table 2.**
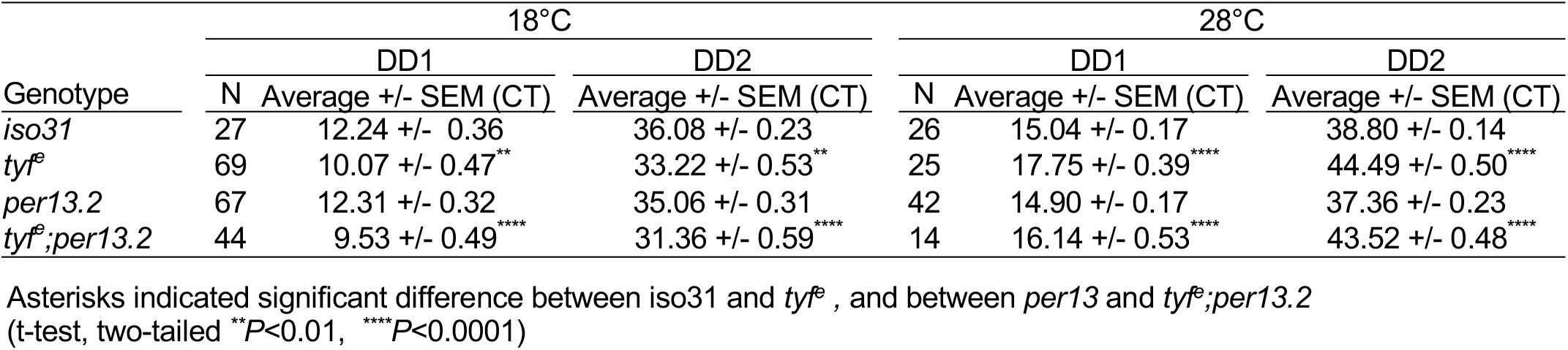
Evening phase offset on the first and second days of DD in *tyf* mutants

To resolve the underlying molecular and cellular basis of these effects, we assessed PER levels on DD1 and DD2 in the sLNv and LNd important for morning, evening, and free-running circadian behavior. At 18°C, we find that in *per13*.*2* PER levels in the sLNv peak at CT24 and fall to a subsequent trough at CT36 while in *tyf; per13*.*2* PER levels start falling from CT18 and trough at CT30 suggesting that *tyf* molecular oscillations are phase advanced (Figure 3). On the other hand, at 28°C in *per13*.*2*, PER levels start peaking at CT18 while peak levels are not reached until CT30-36 suggesting a phase delay consistent with the lengthened period (Figure 3). We observe similar results in the LNd where levels are falling earlier at 18°C with a trough at CT30 (*tyf; per13*.*2*) instead of CT36 (*per13*.*2*) and later peak (CT30; *tyf; per13*.*2* v. CT24; *per13*.*2*) at 28°C consistent with period changes observed (Figure 3, Figure S1). The most robust effects on PER are on peak levels with reductions in peak PER levels in the LNd at both 18°C and 28°C, consistent with a role for *tyf* in activating translation of PER (Figure 3, Figure S1). Notably, effects on the period determining sLNv show significant temperature dependence. Peak PER levels are not significantly changed at 18°C (per13.2 at CT24 v. tyf;per13.2 at CT6, p=0.74) but are at 28°C (per13.2 at CT18 v. tyf;per13.2 at CT36, p<0.05; Figure 3. Figure S1). Thus, *tyf* effects on peak PER levels are temperature dependent, most evident at 28°C versus 18°C which correlates with stronger period effects at 28°C. These results are consistent with a temperature-dependent effect of *tyf* on PER translation.

**Figure 3.**
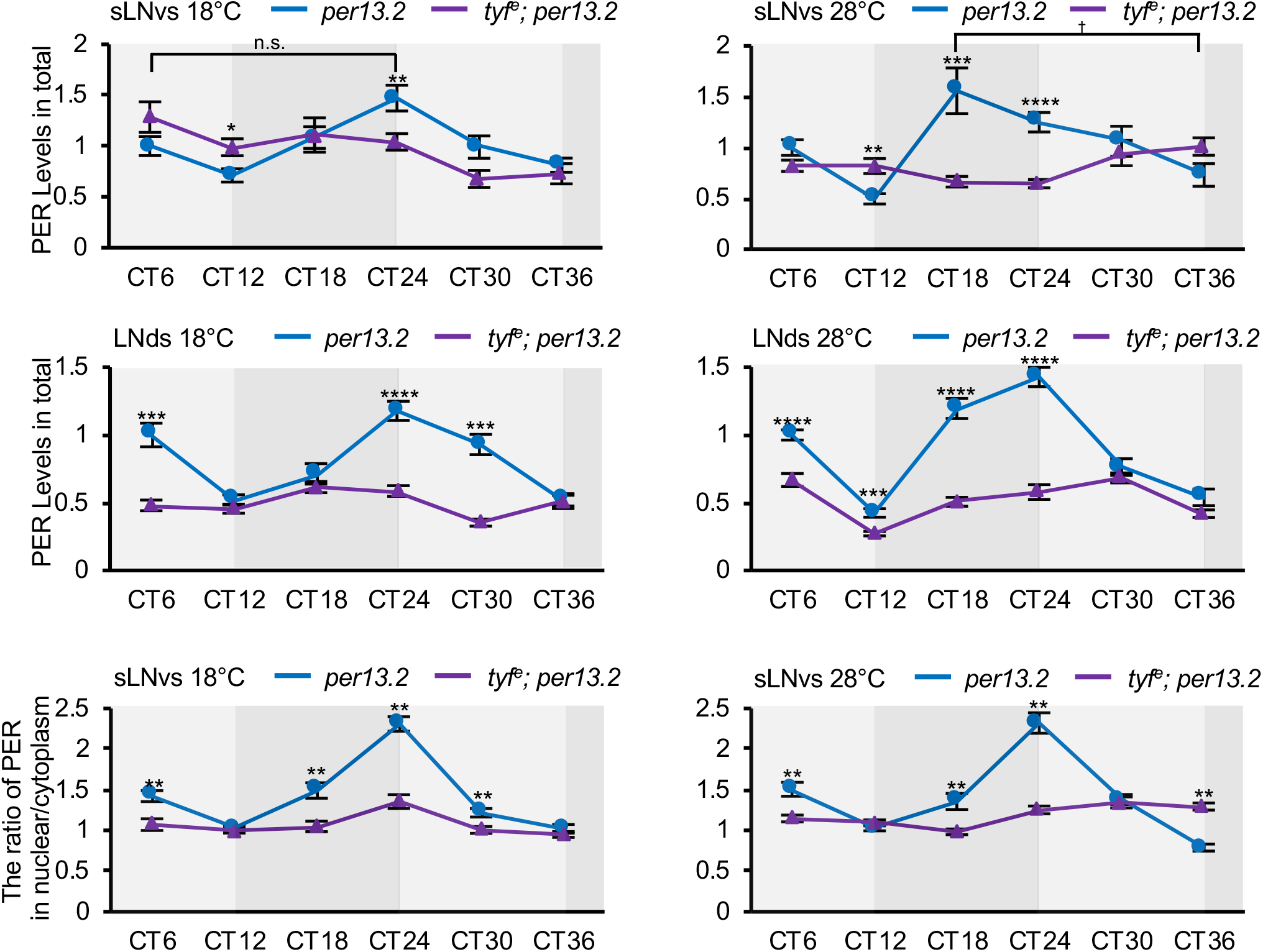
The amount of PER and the peak phase show a temperature dependency in *tyf* mutants. Abundances of PER in sLNvs and LNds were quantified and normalized to the values of *per13*.*2* at CT6 under each temperature condition. The ratio of PER in nuclear and cytoplasm were calculated from the signals detected in each area. Blue and purple lines represent *per13*.*2* and *tyf*^*e*^; *per13*.*2*, respectively. Data represents average ± SEM (*n* = 5 to 20). Asterisks indicate statistically significant differences between *per13*.*2* and *tyf*^*e*^; *per13*.*2* at each time point (t-test, two-tailed, ^*^*P* < 0.05, ^**^*P* < 0.01, ^***^*P* < 0.001 and ^****^*P* < 0.0001). Peak levels of PER show no differences at 18°C indicated as n.s. (*per13*.*2* at CT24 vs *tyf*^*e*^; *per13*.*2* at CT6, t-test, two-tailed, *P* = 0.74). A dagger indicates significant differences of peak levels of PER at 28°C (*per13*.*2* at CT18 vs *tyf*^*e*^; *per13*.*2* at 28°C at CT36, t-test, two-tailed, *P* < 0.05). CT indicates circadian time.

We also examine the PER nuclear-cytoplasmic ratio in the sLNv (Figure 3). Here we found that at 18°C, the N/C ratio peaked at CT24 in both *per13*.*2* and *tyf; per13*.*2* flies, although the rhythm appears blunted in the latter. On the other hand, the rhythm was delayed at 28°C with a broad peak in *tyf; per13*.*2* flies. Thus, we observe molecular changes in the sLNv and LNd that roughly parallel the period changes with an especially strong effect on PER levels in the LNd.

We and others have shown that TYF controls PER translation via a multiprotein complex including ATX2 and LSM12. To independently validate a role for this complex, we assessed temperature compensation in *Lsm12* mutants comparing *Lsm12*^*Δ6*^ to *Lsm12*^*Δ2*^ wild-type controls (Figure 4 and 5, Table 3). *Lsm12* mutants are more rhythmic than *tyf* mutants(23) and thus, we could assess them without additional transgenic *per* copies. As we found for *tyf*, we find that period significantly lengthens in *Lsm12*^*Δ6*^ mutants (Q_10_=0.936 v. 0.983 for the wild-type *Lsm12*^*Δ2*^), although to a lesser degree than *tyf* mutants. To see if PER is rate limiting for this phenotype, we also tested *Lsm12* mutants with *per13*.*2* and found that temperature compensation (Q_10_=0.97) was comparable to the *Lsm12*^*Δ2*^ control flies without *per13*.*2* (Q_10_=0.98) suggesting PER levels are crucial for temperature compensation. To reveal the molecular basis of these effects, we assessed PER levels in the sLNv in *Lsm12* mutants at 18° and 28°C. Consistent with a role in activating PER translation we observed reductions in peak PER levels (ZT0) regardless of temperature (Figure 6A,B).

**Figure 4.**
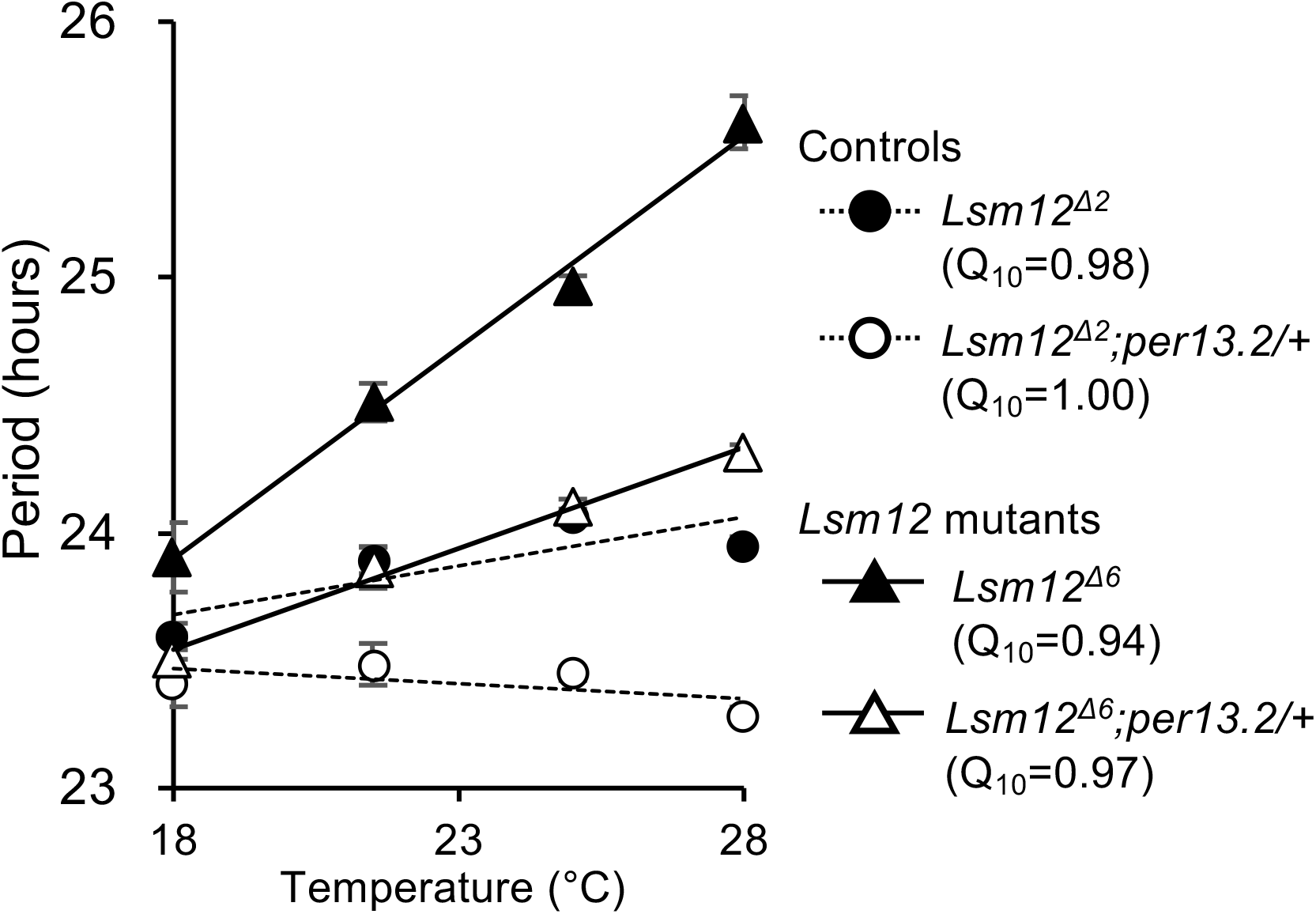
Lack of *Lsm12* shows temperature overcompensation. Averaged periods of the free-running periods under four different temperatures were calculated from the flies showing power - significance values (P-S) over 10 at the corresponding temperatures. Data represents average +/-SEM. The dashed and solid lines represent the regression lines of controls and the *Lsm12* mutants, respectively. *Lsm12*^*Δ6*^ line is a null mutant of *Lsm*12 and *Lsm12*^*Δ2*^ line is the revertant. Sample sizes are indicated in Table 3.

**Figure 5.**
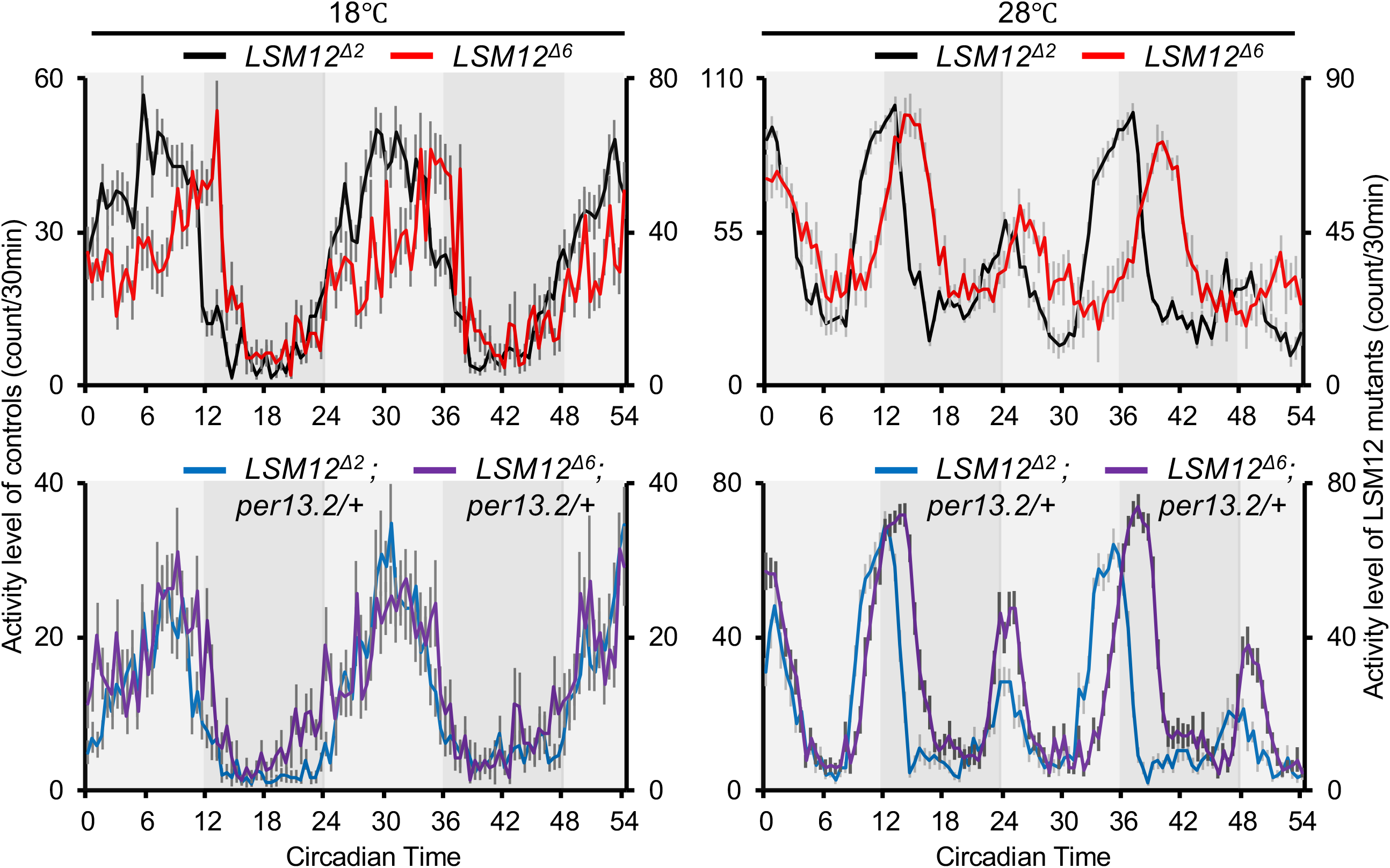
Behavioral period of a *Lsm12* mutant becomes longer under high temperature than low temperature under the first two days of constant darkness. Data represents averaged activity levels +/-SEM. First axis indicates activity level of controls, and the second axis is that of *Lsm12* mutants or *Lsm12* mutants with *per* genomic rescue. Color codes and sample sizes are as follows. Black: *Lsm12*^*Δ2*^ (n=77 and 75 at 18°C and 28°C, respectively). Red: *Lsm12*^*Δ6*^ (n=36 and 39 at 18°C and 28°C, respectively). Blue: *Lsm12*^*Δ2*^*;per13*.*2/+* (n=71 and 72 at 18°C and 28°C, respectively). Purple: *Lsm12*^*Δ6*^*;per13*.*2/+* (n=49 and 50 at 18°C and 28°C, respectively).

**Table 3:** Q10 of behavioral period in *Lsm12* mutants

| Genotype | Temperature<br>(°C) | N | Power +/- SEM | P-S>10 |  |  |  |  |
| --- | --- | --- | --- | --- | --- | --- | --- | --- |
|  |  |  |  | N | Period +/- SEM | Q10 | 2.5%<br>CI | 97.5%<br>CI |
| *Lsm12<sup>Δ2</sup>* | 18 | 77 | 66 +/- 4 | 75 | 23.6 +/- 0.1 | 0.98 | 0.98 | 0.99 |
|  | 21.5 | 56 | 69 +/- 4 | 55 | 23.9 +/- 0.1<sup>a</sup> |  |  |  |
|  | 25 | 41 | 100 +/- 5<sup>a,b</sup> | 40 | 24.1 +/- 0.0<sup>a</sup> |  |  |  |
|  | 28 | 65 | 89 +/- 4<sup>a,b</sup> | 64 | 24.0 +/- 0.0<sup>a</sup> |  |  |  |
| *Lsm12<sup>Δ6</sup>* | 18 | 34 | 45 +/- 5 | 32 | 23.9 +/- 0.2 | 0.94* | 0.93 | 0.95 |
|  | 21.5 | 52 | 60 +/- 5 | 47 | 24.5 +/- 0.1<sup>a</sup> |  |  |  |
|  | 25 | 93 | 49 +/- 4 | 76 | 25.0 +/- 0.0<sup>a,b</sup> |  |  |  |
|  | 28 | 36 | 61 +/- 5 | 33 | 25.6 +/- 0.1<sup>a,b,c</sup> |  |  |  |
| *Lsm12<sup>Δ2</sup>;per13.2/+* | 18 | 70 | 37 +/- 3 | 63 | 23.4 +/- 0.1 | 1.00 | 1.00 | 1.01 |
|  | 21.5 | 68 | 56 +/- 4<sup>a</sup> | 63 | 23.5 +/- 0.1 |  |  |  |
|  | 25 | 67 | 78 +/- 4<sup>a,b</sup> | 67 | 23.4 +/- 0.0 |  |  |  |
|  | 28 | 70 | 96 +/- 4<sup>a,b,c</sup> | 70 | 23.3 +/- 0.0 |  |  |  |
| *Lsm12<sup>Δ6</sup>;per13.2/+* | 18 | 49 | 49 +/- 4 | 45 | 23.5 +/- 0.1 | 0.97* | 0.96 | 0.97 |
|  | 21.5 | 44 | 106 +/- 6<sup>a</sup> | 44 | 23.9 +/- 0.0<sup>a</sup> |  |  |  |
|  | 25 | 44 | 110 +/- 6<sup>a</sup> | 43 | 24.1 +/- 0.0<sup>a,b</sup> |  |  |  |
|  | 28 | 49 | 114 +/- 5<sup>a</sup> | 48 | 24.3 +/- 0.0<sup>a,b,c</sup> |  |  |  |

Statistical significance determined using one-way ANOVA followed Tukey's multiple comparison test. a, b, and c indicate significantly difference with 18°C, 21.5°C and 25°C, respectively. Asterisks show statistically difference between *Lsm12<sup>Δ2</sup>* and *Lsm12<sup>Δ6</sup>*, and *Lsm12<sup>Δ2</sup>;per13.2/+* and *Lsm12<sup>Δ6</sup>;per13.2/+*. CI indicates confidence intervals.

**Table 4.** Evening phase offset on the first and second days of DD in *Lsm12* mutants

| Genotype | 18°C |  |  |  | 28°C |  |  |
| --- | --- | --- | --- | --- | --- | --- | --- |
|  | DD1 |  | DD2 |  | DD1 |  | DD2 |
|  | N | Average +/- SEM (CT) | Average +/- SEM (CT) |  | N | Average +/- SEM (CT) | Average +/- SEM (CT) |
| *Lsm12<sup>Δ2</sup>* | 77 | 12.02 +/- 0.20 | 35.91 +/- 0.19 |  | 75 | 15.47 +/- 0.14 | 39.21 +/- 0.18 |
| *Lsm12<sup>Δ6</sup>* | 36 | 13.76 +/- 0.35**** | 36.98 +/- 0.49* |  | 39 | 17.47 +/- 0.23**** | 42.84 +/- 0.30**** |
| *Lsm12<sup>Δ2</sup>;per13.2/+* | 71 | 12.73 +/- 0.21 | 36.62 +/- 0.28 |  | 72 | 14.79 +/- 0.14 | 37.91 +/- 0.18 |
| *Lsm12<sup>Δ6</sup>;per13.2/+* | 49 | 12.92 +/- 0.26 | 36.02 +/- 0.31 |  | 50 | 16.38 +/- 0.17**** | 40.46 +/- 0.21**** |

Asterisks indicated significant difference between *Lsm12<sup>Δ2</sup>* and *Lsm12<sup>Δ6</sup>*, and between *Lsm12<sup>Δ2</sup>;per13.2/+* and *Lsm12<sup>Δ6</sup>;per13.2/+*. (t-test, two-tailed \**P*<0.05, \*\**P*<0.01, \*\*\*\**P*<0.0001)

**Figure 6.**
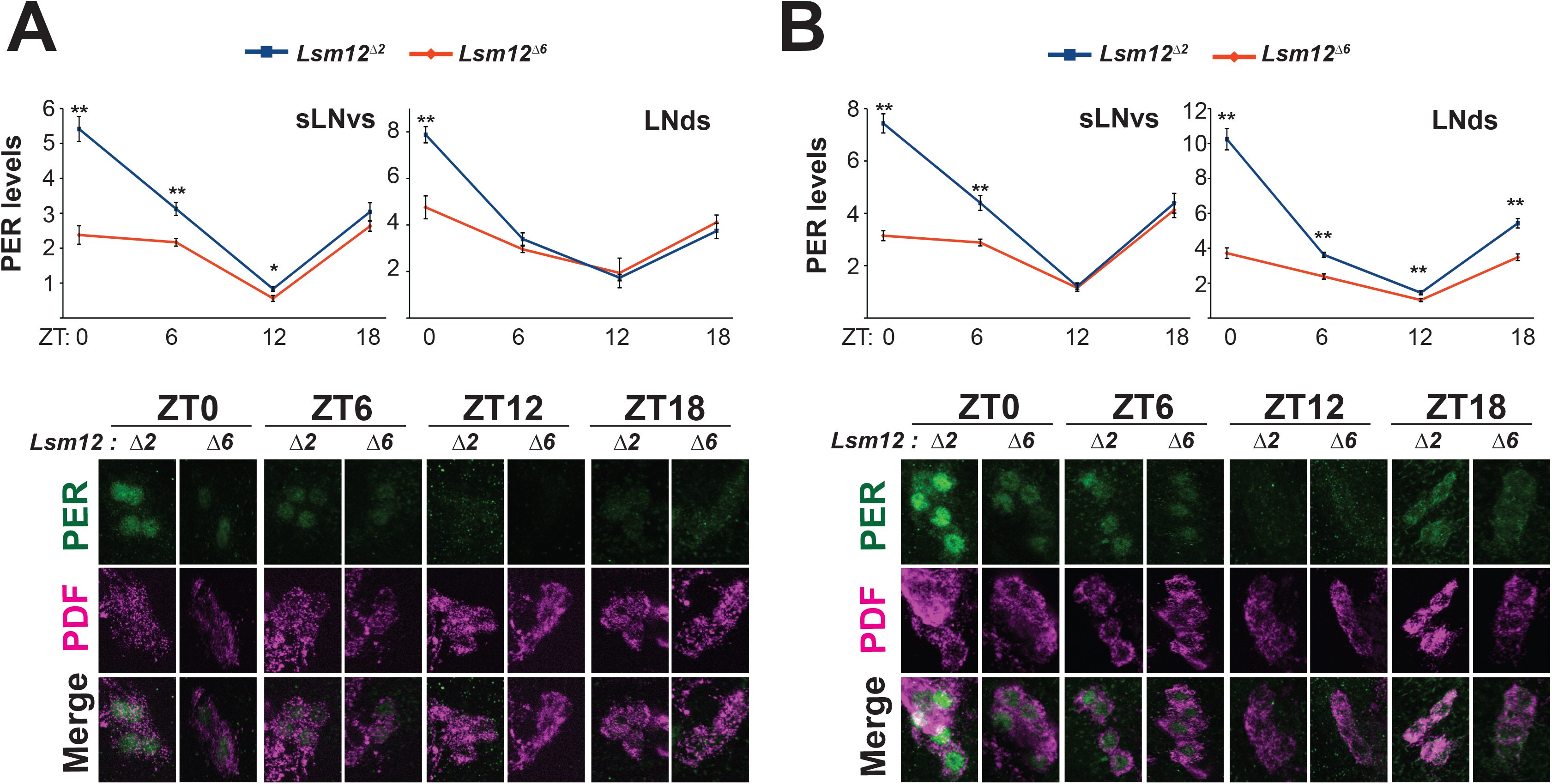
PER levels are dampened in clock neurons of a *Lsm12* mutant. At 18°C (A) and 29°C (B), *Lsm12* mutant flies (lsm12Δ6) exhibit lower PER levels in sLNv’s and LNd’s compared to control flies (*Lsm12*^*Δ2*^). PER levels were quantified from individual pacemaker neurons during LD at four time-points. ZT indicates Zeitgeber time. For normalization the average PER intensity of *Lsm12*^*Δ6*^ at ZT0 was set to 1. Data represent mean ± SEM (on average n=45 per time-point). Statistical comparisons made using Student’s t test (*\*P* < 0.05 and \*\**P* < 0.001). Lower panels: Representative confocal images of *Lsm12*^*Δ2*^ and *Lsm12*^*Δ6*^ sLNv’s at 18°C (A) and 29°C (B).

## Discussion

We report here a genetic perturbation which substantially alters temperature compensation of circadian rhythms in an animal without unambiguously altering the Q_10_ of a gene or protein. Previously described genetic perturbations that alter temperature compensation in animals have or may have altered the Q_10_ of the encoded protein. Even in the case of pharmacological manipulations that alter temperature compensation, direct measures indicate inhibitors/antagonists can also significantly perturb the Q_10_ of their targets(35) or the Q_10_ has not been independently measured(31). In other cases, some gene dosage mutants in fungi that perturb temperature compensation actually exhibit phenotypes with *better* temperature compensation, i.e., Q_10_s closer to 1(5). Here we use two independent null/strongly hypomorphic alleles in combination with or without additional doses of genomic *per* transgenes to demonstrate robust alterations in temperature compensation. As these genetic manipulations should not alter the Q_10_ of their encoded proteins, i.e., TYF and LSM12, they demonstrate their specific role in temperature compensation. This is in contrast to mutants that alter temperature compensation by inducing a temperature sensitive allele in a rate-limiting period determining clock component.

We also demonstrate such temperature compensation phenotypes in an intact animal. While some features of temperature compensation are evident in vitro(7), in unicellular organisms(5) and isolated cells in culture(31), the most robust temperature compensation (i.e., Q10<1.1 and >0.9) is evident in circadian rhythms in multicellular organisms with intact circadian clock networks(13), highlighting a role for pacemaker coupling for the most precise temperature compensation. Our finding of disrupted temperature compensation in animal circadian behavior suggests an important role in this most precise form of temperature compensation.

Our work highlights a specific role of PER translation in temperature compensation. PER translation has been identified as a key regulatory node for robust circadian rhythms driven by a posttranscriptional protein complex including TYF and LSM12. We identified two components of this complex each of which impacts temperature compensation via genetic loss-of-function providing compelling evidence for this complex. Indeed, we can partially rescue these phenotypes in the case of LSM12 by expressing additional copies of PER further underscoring the importance of PER to these phenotypes. Our results experimentally validate proposed translation-based models for temperature compensation (17, 36). In this model, temperature-dependent PER is part of a discrete mechanism that maintains circadian period length. When this mechanism is lost, period length becomes much more temperature sensitive. One intriguing possibility is the idea that *per* mRNA exhibits temperature dependent changes in secondary structure which in turn regulate PER translation. Such structures have been termed RNA thermometers. An alternative is a potential link between the heat shock pathway implicated in mammalian temperature compensation(12) and PER synthesis. Heat shock can rapidly reduce PER levels but this appears to be independent of HSF in *Drosophila*(37). Future studies examining PER metabolism as a function of temperature will be of interest to further examine these models.

Although PER translation appears to be a key temperature compensation mechanism, flies lacking either TYF or LSM12 actually display hyper-compensated rhythms, i.e., they slow down (longer period) with increasing temperature. Thus, there remains to be determined what other mechanisms contribute to temperature compensation, such as temperature-dependent phosphorylation and degradation(31), that may act to slow the clock with temperature in opposition to PER translation. Genetic perturbations that do not themselves alter the Q_10_ of the perturbed gene, e.g., changes in gene dosage, as we have demonstrated here, will be essential to establish these mechanisms. We hypothesize that the multiple regulatory layers of the clock may have evolved to not only drive free running rhythmicity but period stability in response to various environmental challenges such as changing temperature.

## Acknowledgements

This research was supported by grants JSPS KAKENHI(18K14749) to T.Q.I., NIH (R01NS106955) to R.A., NSF (DMS-176442) to R.A/R.B., Simons Foundation (SFARI 597491-RWC) to R.A./R.B., and DARPA (D12AP00023) to R.A. The content of the information does not necessarily reflect the position or the policy of the Government, and no official endorsement should be inferred. We thank C. Hong for constructive discussion and C. Lim for *Lsm12* strains.

## Materials and Methods

### *Drosophila* Strains

All flies were kept on standard cornmeal food under 12-h light:12-h dark (LD) cycles at 25 °C. *iso31* (*w*^*1118*^; BL5905) was acquired from Bloomington *Drosophila* Stock center. A null mutant of *tyf* (*tyf*^*e*^) was previously described(25). The transformant containing a 13.2-kb genomic fragment of *period* locus (*per*^*0*^; *per13*.*2*)(38), and a LSM12 mutant (*LSM*^*Δ2*^) and the revertant (*LSM*^*Δ6*^)(23) were kindly provided by the Rosbash lab and by the Lim lab, respectively. *per13*.*2, tyf*^*e*^; *per13*.*2/+, tyf*^*e*^; *per13*.*2, LSM*^*Δ2*^; *per13*.*2/+* and *LSM*^*Δ6*^; *per13*.*2/+* were established by the standard crossing scheme using the strains described above. All data shown in this paper was obtained from adult male flies.

### Locomotor Analysis

Recording and analysis of *Drosophila* locomotor behavior using the Drosophila Activity Monitoring (DAM) system (*Drosophila* activity monitor, Trikinetics) were described previously(39, 40). Briefly, individual progeny raised at 25 °C were loaded into tubes containing 1% agar, 5% sucrose food. The DAM system monitored and recorded their behavior under five LD cycles followed by seven days under constant darkness at each indicated temperature condition (18°C, 21.5°C, 25°C and 28°C). Unless otherwise noted, period was calculated from χ^2^ periodogram using Clocklab (Actimectrics) from the individual fly shown “Power-Significant” (P-S) value ≥ 40, and averaged in each genotype. Evening offset under DD1 and DD2 were calculated from the mean of their peak and trough phase manually selected from the activity level defined as number of beam crossings per 30min in individual flies.

### Q_10_ Statistical Methods

*Q*10 values for circadian oscillators are according to the usual definitiion

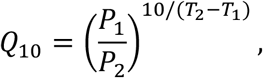

Where *P*_1_ and *P*_2_ are the periods at temperatures *T*_1_ and *T*_2_ respectively. Simple algebraic manipulation yields

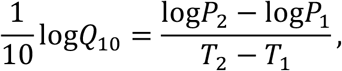

the right-hand-side of which may be simply estimated from a linear regression of the data. *Q*_10_ estimates and confidence intervals are thus obtained as *Q*_10_ = exp(10. *β*_1),_ where *β*_1_ is the estimated slope of the regression line log(*P*) = *β*_0_ + *β*_1_*T*. Statistical significance of differential *Q*_10_ between genotypes is assessed via the significance of an interaction term *β*_2_ in the model log(*P*) = *β*_0_ + *β*_1_*T* + *β*_2_*X*_*genotype*_*T*.

### Immunofluorescence

Flies aged ≥5 d were transferred into DD conditions after the 2 days of entrainment or collected in LD as indicated. Brains of each genotype were dissected and immunostained with antibodies against PER as follows. Each fly was sampled at a certain time point and their brains were dissected in PBS within 10 minutes. The dissected brains were fixed in 3.7% formalin solution (diluted from 37% formalin solution, Sigma-Aldrich) for 30 minutes. 0.3% PBSTx (PBS with 0.3% Triton-X) washed brains for 4 times followed by the incubation in a primary antibody diluted in 0.3% PBSTx with 5% normal goat serum at 4°C overnight. Brains were washed for 4 times with 0.3% PBSTx after primary antibody incubation. Secondary antibodies were diluted in 0.3% PBSTx with 5% normal goat serum and incubated with brains at 4°C overnight. The dilution of primary and secondary antibodies was as follow: mouse anti-PDF (1:800, DSHB), rabbit anti-PER (1:8000, from the Rosbash Lab), anti-mouse Alexa594 (1:800, invitrogen), anti-rabbit Alexa488 (1:800, invitrogen). After mounting all brains with VECTASHIELD (Vector Labs), each sample was kept under 4°C until observation.

### Immunofluorescence Quantification

Immunostained brain samples were imaged by a Nikon C2 confocal microscope. The processing of data and signal quantification were performed using NIH ImageJ/Fiji. Confocal images of PDF positive sLNvs, and LNds were z-stacked with their maximum intensity of signals. Quantification of PER signal was performed as described previously(41). PER signal intensity above background was calculated using the formula: I = (S − B)/B, where *S* is the fluorescence intensity, and *B* is the average intensity of the region adjacent to the cells. The signal values obtained from a single cell were averaged within the corresponding hemisphere, then were averaged by the number of imaged hemispheres for *tyf* mutant experiments. The signal values were finally normalized by the signal of a certain time point in control strain. In Lsm12 mutant experiments, we averaged intensity from individual cells from different brains(42).

## Notes

### Competing Interest Statement

The authors have declared no competing interest.

