## Supplemental Figure 1 for "*period* Translation as a Core Mechanism Controlling Temperature Compensation in an Animal Circadian Clock"

### Supporting Figure Legends

#### Figure S1. Temperature Dependent PER Oscillations in *tyf* Mutants

Representative images of PER in sLNvs and LNds under each temperature condition.

PER and PDF signals are shown as green and magenta, respectively.
